# DNA from fecal immunochemical test can replace stool for microbiota-based colorectal cancer screening

**DOI:** 10.1101/048389

**Authors:** Nielson T. Baxter, Charles C. Koumpouras, Mary A.M. Rogers, Mack T. Ruffin, Patrick D. Schloss

**Affiliations:** Department of Microbiology and Immunology, University of Michigan, Ann Arbor, Michigan.; Department of Internal Medicine, University of Michigan, Ann Arbor, Michigan.; Department of Family Medicine, University of Michigan, Ann Arbor, Michigan.

**Keywords:** colorectal cancer, gut microbiome, microbiota, fecal immunochemical test, random forest

## Abstract

**Background:** There is a significant demand for colorectal cancer (CRC) screening methods that are noninvasive, inexpensive, and capable of accurately detecting early stage tumors. It has been shown that models based on the gut microbiota can complement the fecal occult blood test and fecal immunochemical test (FIT). However, a barrier to microbiota-based screening is the need to collect and store a patient’s stool sample.

**Methods:** Using stool samples collected from 404 patients we tested whether the residual buffer containing resuspended feces in FIT cartridges could be used in place of intact stool samples.

**Results:** We found that the bacterial DNA isolated from FIT cartridges largely recapitulated the community structure and membership of patients’ stool microbiota and that the abundance of bacteria associated with CRC were conserved. We also found that models for detecting CRC that were generated using bacterial abundances from FIT cartridges were equally predictive as models generated using bacterial abundances from stool.

**Conclusions:** These findings demonstrate the potential for using residual buffer from FIT cartridges in place of stool for microbiota-based screening for CRC. This may reduce the need to collect and process separate stool samples and may facilitate combining FIT and microbiota-based biomarkers into a single test. Additionally, FIT cartridges could constitute a novel data source for studying the role of the microbiome in cancer and other diseases.

## Background

Although colorectal cancer (CRC) mortality has declined in recent decades, it remains the second leading cause of death among cancers in the United States [1]. Early detection of CRC is critical since patients whose tumors are detected at an early stage have a greater than 90% chance of survival [1]. However more than a third of individuals for whom screening is recommended do not adhere to screening guidelines [2]. The high cost and invasive nature of procedures, such as colonoscopy and sigmoidoscopy are barriers for many people [3, 4]. Unfortunately non-invasive tests, such as the guaiac fecal occult blood test (gFOBT), fecal immunochemical test (FIT), and the multitarget DNA test fail to reliably detect adenomas [5, 6] (e.g., sensitivity for nonadvanced adenomas is 7.6% for FIT and 17.2% for the DNA test). Thus, there is a need for novel non-invasive screening methods with improved sensitivity for early stage colonic lesions.

Several studies have demonstrated the potential for the gut microbiota to be used to detect CRC [7–10]. Moreover, we and others have shown that combining microbiota-analysis with conventional diagnostics, like gFOBT and FIT, can significantly improve the detection of colonic lesions over either method by itself [7, 8, 10]. One limitation of microbiota-based CRC screening is the need to collect and process separate stool samples for microbiota characterization. Given the widespread use of FIT to collect specimens for screening, the ability to use the same sample for microbiota characterization could make processing more efficient and less expensive. We hypothesized that the small amount of fecal material contained in FIT sampling cartridges was sufficient to perform both hemoglobin quantification and microbiota characterization. To test this hypothesis, we isolated bacterial DNA from the residual buffer of OC-Auto^®^ FIT cartridges (Polymedco Inc.) that had already been used for quantifying fecal hemoglobin concentrations. We then compared the bacterial composition of the FIT cartridge to that of DNA isolated directly from a patient’s stool sample and assessed the ability of FIT cartridge-derived DNA to be used for microbiota-based CRC screening.

## Materials and Methods

### Study Design / Diagnoses / Stool Collection

Stool samples were obtained through the Great Lakes-New England Early Detection Research Network. Patients were asymptomatic, at least 18 years old, willing to sign informed consent, able to tolerate removal of 58 mL of blood, and willing to collect a stool sample. Patient age at the time of enrollment ranged from 29 to 89 with a median of 60 years. Patients were excluded if they had undergone surgery, radiation, or chemotherapy for current CRC prior to baseline samples or had inflammatory bowel disease, known hereditary non-polyposis CRC, or familial adenomatous polyposis. Patient diagnoses were determined by colonoscopic examination and histopathological review of any biopsies taken. Colonoscopies were performed and fecal samples were collected in four locations: Toronto (Ontario, Canada), Boston (Massachusetts, USA), Houston (Texas, USA), and Ann Arbor (Michigan, USA). Stool samples were packed in ice, shipped to a processing center via next day delivery and stored at ‐80°C. Fecal material for FIT was collected from frozen stool aliquots using OC-Auto^®^ FIT sampling bottles (Polymedco Inc.), processed using an OC-Auto Micro 80 automated system (Polymedco Inc.), and stored at ‐20C. The University of Michigan Institutional Review Board approved this study, and all subjects provided informed consent.

### 16S rRNA gene sequencing

Processed FIT samples were thawed, and 100 μl of buffer were withdrawn by pipette for DNA extraction. DNA was isolated from FIT samples or matching stool samples using the PowerSoil-htp 96 Well Soil DNA isolation kit (MO BIO Laboratories) and an epMotion 5075 automated pipetting system (Eppendorf). The V4 region of the bacterial 16S rRNA gene was amplified using custom barcoded primers and sequenced as described previously using an Illumina MiSeq sequencer [11]. The 16S rRNA gene sequences were curated using the mothur software package, as described previously [11, 12]. Curated sequences were clustered into operational taxonomic units (OTUs) using a 97% similarity cutoff with the average neighbor clustering algorithm. Sequences were classified using a naive Bayesian classifier trained against a 16S rRNA gene training set provided by the Ribosomal Database Project (RDP) [13]. Species-level classifications for OTUs of interest were determined by using blastn to compare the predominant sequence within each OTU to the NCBI 16S rRNA database. The putative species was only reported for OTUs with greater than 99% sequence identity to a single species in the database; otherwise the consensus RDP classification was used.

### Statistical Methods

All statistical analyses were performed using R (v.3.2.0). Random forest models were generated using the AUC-RF algorithm for feature reduction and maximizing model performance [14]. The most predictive OTUs were determined based on mean decrease in accuracy when removed from the model. The area under the curve (AUC) of receiver operator characteristic (ROC) curves were compared using the method described by DeLong et al. [15] as implemented in the pROC R package [16].

## Results

DNA was isolated and 16S rRNA gene sequencing was performed on stool aliquots and the residual buffer of paired OC-Auto^®^ FIT sampling cartridges from 404 patients. Among these patients, 101 had CRC, 162 had adenomas, and 141 had no colonic lesions. First, we tested whether the bacterial community profiles from FIT cartridges recapitulated their stool counterparts. First, we compared the number of OTUs shared within FIT/stool pairs from the same patient to the number of OTUs shared between patients (Fig. 1A). FIT cartridges and stool from the same patient (red line) had significantly more bacterial populations in common than those taken from different patients (p<0.001, two-sample Kolmogorov-Smirnov test), indicating that community membership was conserved within patients across stool and FIT cartridges. Second, we calculated the similarity in community structure between samples using 1-thetaYC index [17]. This metric compares the presence or absence of bacterial populations and their relative abundance. The bacterial community structure of stool and FIT samples from the same patient (red line) were significantly more similar to each other than to stool or FIT from other patients (Fig. 1B, p<0.001). Finally, we used a Mantel test to determine whether the patient-to-patient thetaYC distances among stool samples were correlated with the patient-to-patient thetaYC distances among FIT cartridges. We found that there was a significant correlation (Mantel test r=0.525, p<0.001), suggesting that the interpatient variation in community structure between the stool samples of patients was conserved in samples from FIT cartridges.

**Figure 1.**
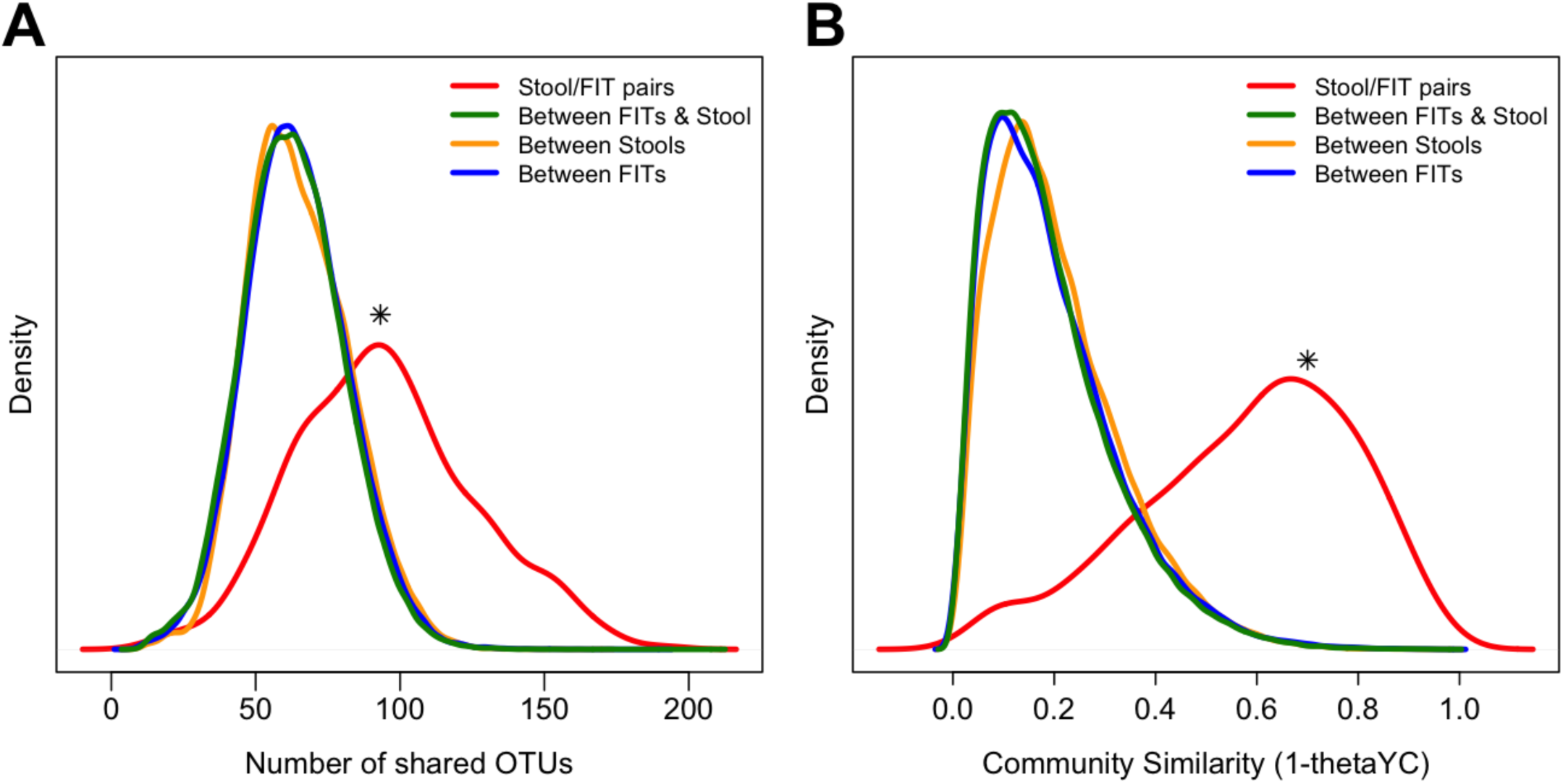
Bacterial community structure from FIT cartridge recapitulates stool. Density plots showing distribution of the number of shared OTUs (A) and community similarity (B) between groups of samples (* p<0.001 two-sample Kolmogorov-Smirnov Test).

Next, we observed a significant correlation between the abundance of each genus in the paired FIT cartridge and stool samples (Fig. 2A, Spearman rho: 0.699, p<0.001). This suggested that the abundance of bacterial genera was conserved. This correlation was especially strong when comparing only the 100 most abundant genera from stool (Spearman rho: 0.886, p<0.001). Several bacterial species have been repeatedly associated with CRC, including *Fusobacterium nucleatum, Porphyromonas asaccharolytica, Peptostreptococcus stomatis*, and *Parvimonas micra* [8–10, 18]. As expected, the abundance of these species in stool was significantly correlated with their abundance in matched FIT cartridges (all p<0.001, Spearman rho ≥0.352)(Fig. 2B). We observed some biases in the abundance of certain taxa. In particular, the genus *Pantoea* was detected in 399 of the 404 FIT cartridges with an average abundance of 2.4%, but was only detected in 1 stool sample. The genus *Helicobacter* was detected in 172 FIT cartridges, but only 10 stool samples. Likewise several genera of *Actinobacteria* were more abundant in stool samples compared to FIT. Notwithstanding these few exceptions, the abundance of the vast majority of genera were well conserved between stool and FIT cartridges. Overall, these findings suggested that that the overall bacterial community structure and the abundance of specific taxa in FIT cartridges and stool were similar.

**Figure 2.**
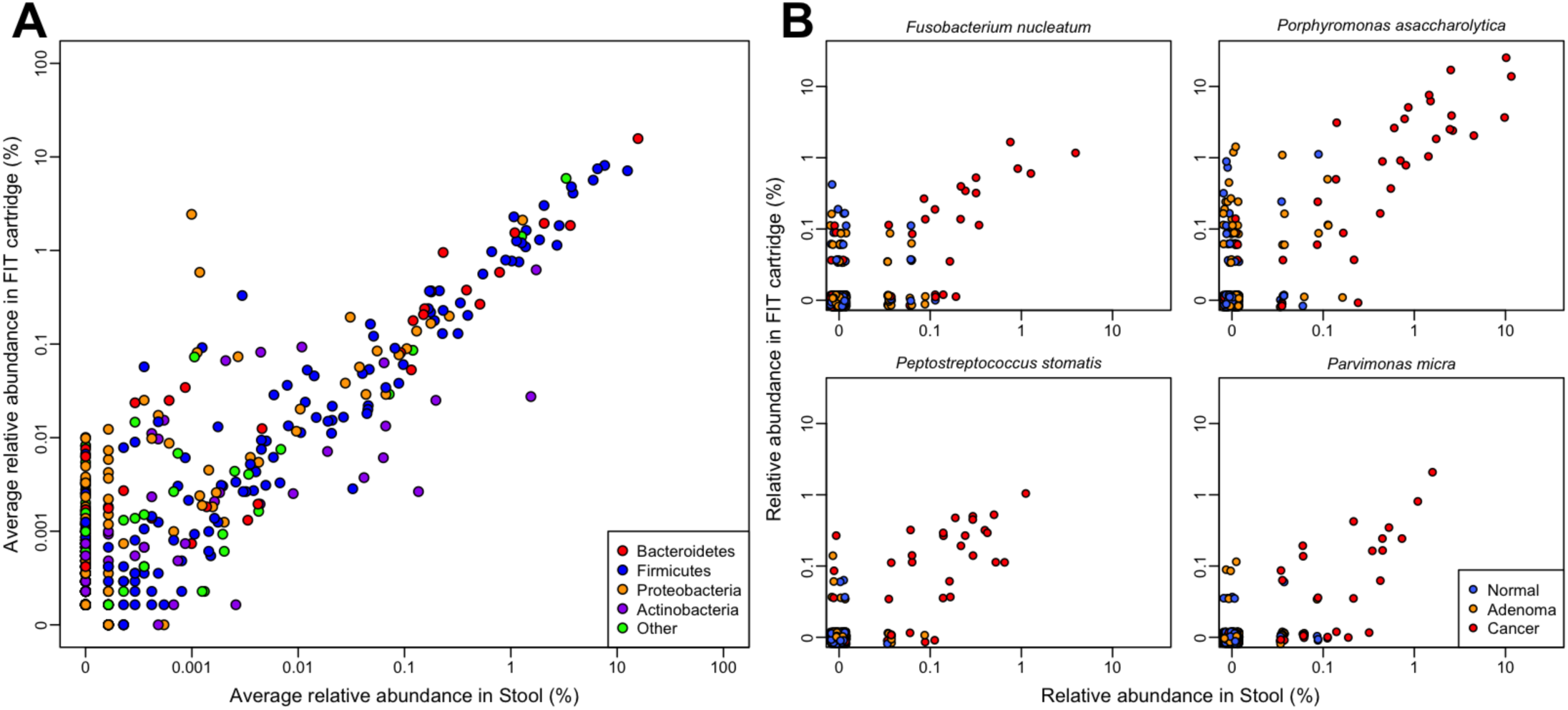
Bacterial populations conserved between stool and FIT cartridge. (A) Scatterplot of the average relative abundance of each bacterial genus in stool and FIT cartridges colored by phylum. (B) Scatterplots of the relative abundances of 4 species frequently associated with CRC. All correlations were greater than 0.35 (all p<0.001).

We tested whether the bacterial relative abundances we observed from FIT cartridges could be used to differentiate healthy patients from those with carcinomas using random forest models as we did previously using intact stool samples [10]. Using DNA from the FIT cartridge, the optimal model utilized 28 OTUs and had an AUC of 0.831 (Fig. 3A). There was not a significant difference in the AUC for this model and the model based on DNA isolated directly from stool, which used 32 OTUs and had an AUC of 0.853 (p=0.41). Furthermore, the probabilities of individuals having lesions were correlated between the models generated using DNA isolated from the FIT cartridges and stool samples (Spearman rho: 0.633, p<0.001, Fig. 3B). We also generated random forest models for differentiating healthy patients from those with any type of lesions (i.e. adenoma or carcinoma). There was not a significant difference in AUC between the stool-based model with 41 OTUs (AUC=0.700) and the FIT cartridge-based model with 41 OTUs (AUC=0.686, p=0.65, Fig. 3C). Again, the probabilities of individuals having lesions according to the two models were significantly correlated (Spearman rho: 0.389, p<0.001 Fig. 3D). These findings demonstrated that models based on bacterial DNA from FIT cartridges were as predictive as models based on DNA isolated directly from stool.

**Figure 3.**
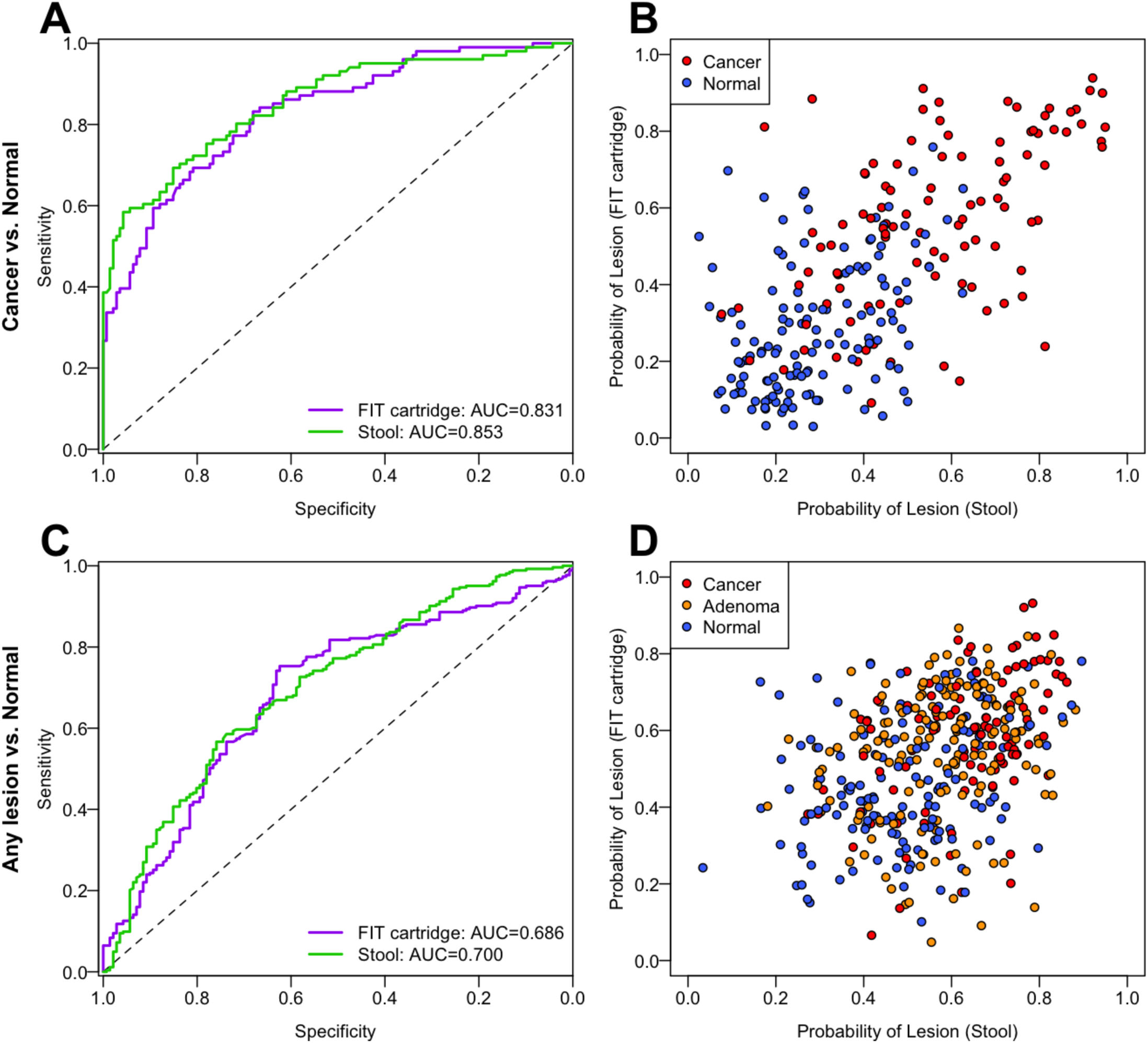
Microbiota-based models from FIT cartridge DNA are as predictive as models from stool. (A) ROC curves for distinguishing healthy patients from those with cancer using microbiota-based random forest models using DNA from FIT cartridges or stool. (B) Probability of having cancer for each patient according to microbiota-based models from A. (C) ROC curves for distinguishing patients with adenomas or carcinomas from healthy patients using microbiota-based random forest models using DNA from FIT cartridges or stool. (D) Probability of having a lesion for each patient based on the models from C.

## Discussion

Bacterial DNA isolated from the residual buffer of FIT cartridges recapitulated the community structure and membership of patients’ stool microbiota. FIT/stool pairs collected from the same patient were significantly more similar to each other than samples from different patients and the inter-patient differences in stool microbiota structure were conserved in FIT cartridge-derived microbiota. More importantly, random forest models generated using bacterial abundances from FIT cartridge-derived and stool-derived DNA were equally predictive for differentiating healthy patients from those with adenomas and carcinomas.

Sinha et al. compared a variety of sampling and storage methods for fecal samples to be used for microbiome analyses [19]. They found reproducible biases according to sampling method and time at ambient temperature. Likewise, we observed biases in the abundance certain bacterial populations in FIT cartridges compared stool. For example, an OTU associated with *Pantoea* was found in 98.8% of FIT cartridge samples and only 0.2% of stool samples. There are several possible explanations for this result. It is possible that because the biomass contained in the FIT cartridges is considerably lower than that in stool, the analysis was more sensitive to contaminants in our reagents or the FIT cartridge [20]. Alternatively, storage conditions could have played a role in biasing the relative abundances of certain genera. The feces in the FIT cartridges spent more time exposed to ambient temperatures in order to be analyzed for hemoglobin concentration. Therefore it is possible that certain bacterial populations, especially aerobes, were able to grow. Considering *Pantoea* is rarely found in human feces and is more commonly found in soil, plant surfaces, and air we suspect that it was a contaminant. Regardless of the source of this and the other suspicious populations, any biases were limited since the random forest feature selection process did not select these populations and did not affect the ability to detect CRC from FIT cartridge-derived DNA.

## Conclusions

This could reduce the need to collect and process separate stool samples, decreasing the cost of screening. It may be possible to use FIT cartridges rather than separate stool samples for future studies on the role of the gut microbiota and cancer. Samples collected from patients who undergo annual FIT screening could be used to monitor temporal changes in a patient’s microbiota, making it possible to detect shifts toward a disease-associated microbiota. Since FIT cartridges are currently used for CRC screening, our findings may facilitate large-scale validations of microbiota-based screening methods.

## Declarations Abbreviations

FIT: fecal immunochemical test
gFOBT: guaic fecal occult blood test
OTU: operational taxonomic unit
AUC: area under the curve
ROC curve: reciever operating characteristic curve

## Ethics approval and consent to participate

The University of Michigan Institutional Review Board approved this study, and all subjects provided informed consent. This study conformed to the guidelines of the Helsinki Declaration.

## Availability of data and materials

Raw fastq files and a MIMARKS file are available through the NCBI Sequence Read Archive [SRP062005]. The data processing steps for going from the raw sequence data to the final manuscript is available at http://www.github.com/SchlossLab/Baxter_FITs_BMCCancer_2016.

## Author Contributions

PDS, MTR, MAMR, and NTB were involved in the conception and design of the study. NTB and CCK performed DNA extractions and 16S rRNA gene sequencing. NTB analyzed the data. All authors interpreted the data. NTB and PDS wrote the manuscript. All authors reviewed and revised the manuscript.

## Acknowledgements

The authors thank the Great Lakes-New England Early Detection Research Network for providing the fecal samples that were used in this study.

